# No-code microbial growth phenotyping with GUIbiont

**DOI:** 10.64898/2026.08.17.745250

**Authors:** Edgar Z. Alvarenga, Edoardo Oltolini, Fernanda Pinheiro

## Abstract

Microbial growth screens generate thousands of curves, but cross-experiment comparison and mapping growth phenotypes to genotypes or environments routinely require custom code. `GUIbiont` is a no-code browser application for quality control, curve fitting, clustering and metadata-linked analysis with machine learning techniques. Interactive sessions export as Julia scripts, allowing users to reproduce or extend browser analyses. Validated across 3,885 *E. coli* deletion strains and 13,608 defined-media curves, `GUIbiont` recovered known auxotrophic and nutrient-dependent phenotypes.

---

Biological questions in microbial growth screens often extend across experimental days, strains and media, requiring comparisons among replicate plates and growth curves. Although many tools automate curve fitting and recent interfaces add interactive quality control [1-6], fitting is only one component of screen-level analysis. Pooling runs under consistent settings, classifying trajectories and relating fitted parameters to metadata typically fall outside these curve-fitting tools and are handled with custom scripts (Supp. Table S1). This can separate interpretation from data collection, making it harder to inspect results as experiments proceed, adapt the analysis and reuse it for new data.

`GUIbiont` addresses this gap with a browser-based application for the analysis of microbial growth data built on `Kinbiont.j1`, a Julia framework for data preprocessing, model-based parameter inference and downstream analysis with interpretable machine learning methods [7]. In developing `GUIbiont`, we extended `Kinbiont.j1` with methods for clustering and made them available through the browser interface. Without writing Julia code, users can import plate-reader files and annotations, inspect raw curves and replicate means, fit individual curves or entire screens, cluster full trajectories into phenotypic classes and relate fitted parameters to genotypes or medium compositions (Fig. 1). For single-curve fitting, batch fitting, and clustering, `GUIbiont` turns user-selected browser settings into downloadable Julia scripts, so that the interactive analyses can be reproduced or extended directly in Julia without being reconstructed from scratch.

**Figure 1.**
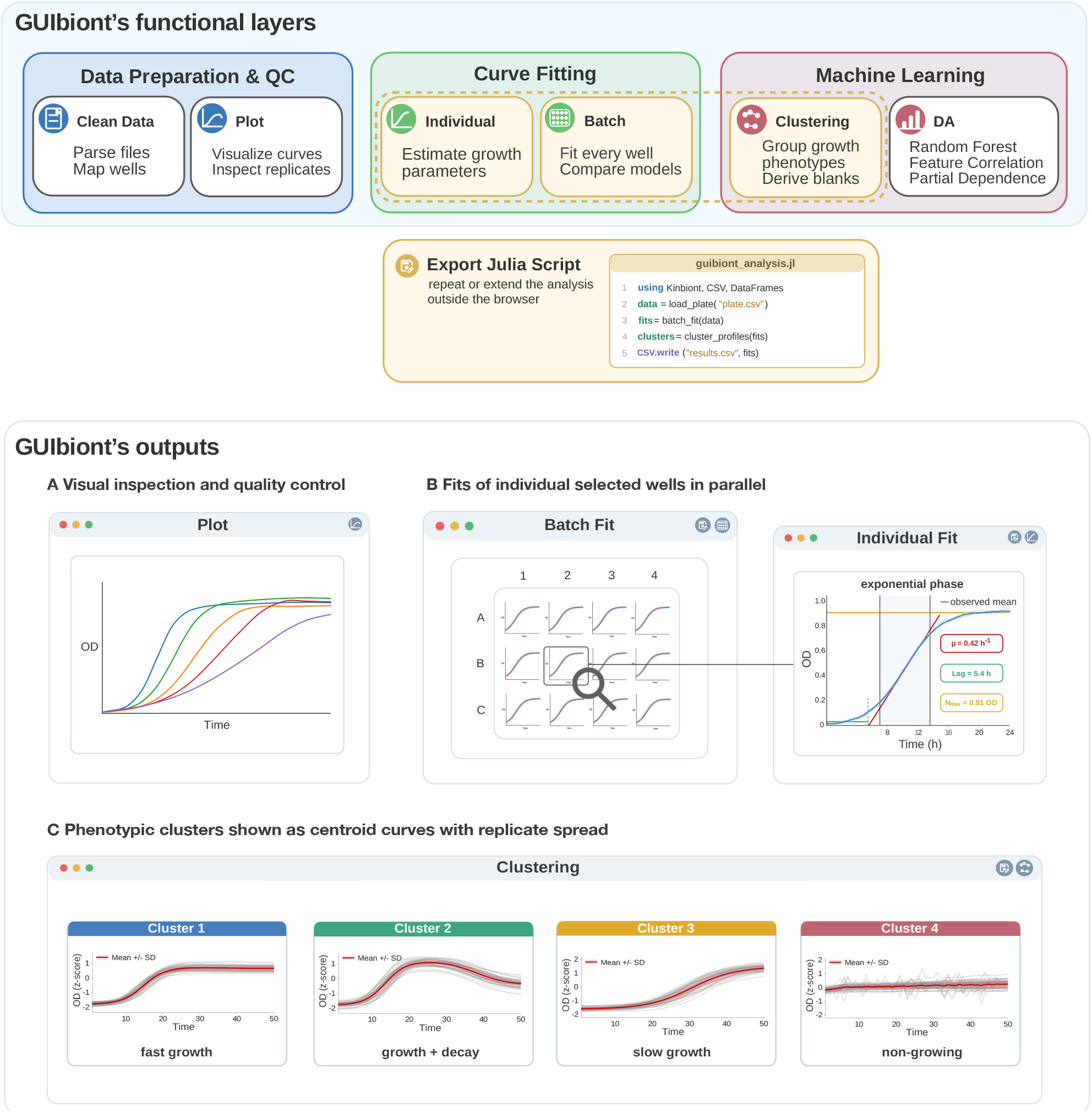
GUIbiont functional layers and representative outputs. *Top:* The browser interface organizes growth-phenotyping analyses into data preparation and quality control, curve fitting and machine learning. *Clean Data* parses plate-reader files and maps wells, while *Plot* supports visual inspection of curves and replicates. *Individual* and *Batch* estimate growth parameters for selected curves or complete screens. *Clustering* groups growth trajectories and supports data-driven blank estimation, while *Down-stream Analysis {DA)* relates fitted parameters to metadata through feature correlations, random forests and partial-dependence analysis. The gold outline marks workflows that can be exported as executable Julia scripts for reproduction or extension outside the browser; the inset illustrates the structure of an exported script. *Bottom:* Representative outputs include **A** curve visualization, **B** parallel fitting of selected wells, an individual fit reporting growth rate *µ*, lag and *N*_max_, and **C** phenotypic clusters displayed as centroid curves with replicate variation. The illustrative cluster labels describe differences in trajectory shape and timing.

We first evaluated `GUIbiont` on a genome-scale screen of 3,909 *E. coli* single-gene deletion strains from the Keio collection, grown in rich LB and M63 minimal media [8]. After aggregation by gene symbol, the screen comprised 3,885 gene-level profiles. Visual inspection of the full dataset in the *Plot* panel showed a subset of curves with no apparent growth in M63 (Fig. 2A), whereas no comparable pattern was observed in LB. The optional constant-curve pre-screen assigned 97 M63 profiles (2.5%) to a separate non-growing class. After separating this class, clustering resolved two dynamic profiles among the growing trajectories in each medium, with centroids differing primarily in the timing of the growth transition (Fig. 2B-C; Methods). Of the 97 non-growing profiles, 82 (84.5%) represented deletions in genes assigned to amino-acid, cofactor or nucleotide biosynthesis pathways, including 49 assigned to amino-acid biosynthesis. The broader biosynthesis-related set and the amino-acid biosynthesis set were enriched 11.8-fold and 19.1-fold, respectively, relative to the complete collection (one-sided Fisher s exact tests, both *P <* 10^−6^; Supplementary Data S2). Thus, the pre-screen recovered a known minimal-medium auxotrophy signature without using genotype annotations to define the non-growing class. Batch fitting with the log-linear sliding-window estimator returned converged estimates of the maximum specific growth rate, *µ*_max_, for 7,767 of the 7,770 gene-medium trajectories in approximately twelve seconds on a six-core workstation (Methods; Supplementary Data S1).

**Figure 2.**
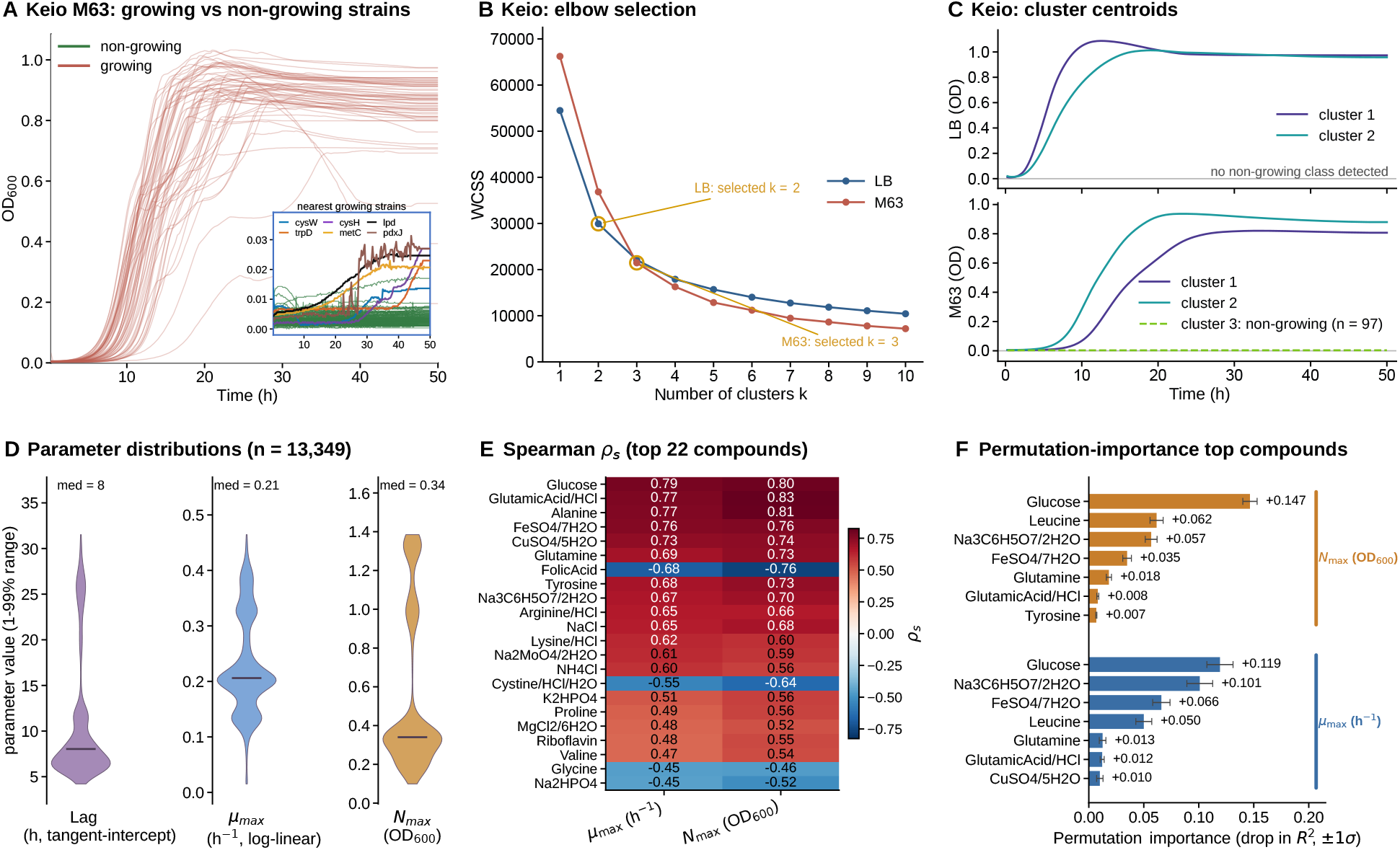
GUIbiont applied to two complementary microbial growth datasets. Top row: *E. coli* Keio collection, comprising 3,909 single-gene deletion strains aggregated into 3,885 gene-level profiles, grown in LB and M63. **A** M63 growth curves for a sample of the 3,788 growing gene-level profiles and all 97 profiles assigned to the non-growing class by the optional pre-screen. The inset shows the non-growing trajectories at low OD together with the six growing profiles closest to the non-growing centroid for comparison; all non-growing trajectories remained below OD_600_ = 0.018 throughout the experiment. **B** WCSS curves used to select *k* = 2 for LB and *k* = 3 for M63, indicated by gold circles. Because the M63 non-growing profiles were separated before clustering, the reported *k* = 3 comprises the non-growing class and two clusters of growing trajectories. **C** Cluster centroids on the raw OD scale. The two growing profiles in each medium differ primarily in the timing of the growth transition. Bottom row: *E. coli* BW25113 grown in 1,029 chemically defined media varying in the concentrations of 44 compounds. **D** Distributions of the maximum specific growth rate *µ*_max_, tangent-intercept lag and saturation OD *N*_max_ estimated with the log-linear sliding-window procedure across the 13,349 fitted curves. Medians and the 1st to 99th percentile ranges are shown. **E** Spearman correlations between compound concentrations and *µ*_max_ or *N*_max_ across the 1,026 media retained after replicate averaging, for the 22 compounds with the strongest associations with *µ*_max_; all values are shown. **F** Random-forest permutation importance for the seven highest-ranking compounds for *µ*_max_ and *N*_max_, reported as the decrease in training-set *R*^2^ after shuffling each compound. Error bars show one standard deviation across ten shuffles.

We next asked how well medium composition predicts growth phenotypes in a fixed genetic background. We analyzed 13,608 growth curves from *E. coli* BW25113 grown in 1,029 chemically defined media varying in the concentrations of 44 compounds [9]. Quality control retained 13,400 curves across 1,026 media (Methods). Log-linear batch fitting returned 13,349 converged estimates of *µ*_max_ and *N*_max_ (Fig. 2D). We then averaged replicates within each medium and focused the compound-level analysis on *µ*_max_ and *N*_max_, as complementary measures of maximum growth rate and saturation optical density (OD). Random forests evaluated by fivefold cross-validation predicted both parameters in held-out media 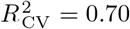 and 0.87, respectively). Lucose had the strongest univariate association with *µ*_max_ (Spearman coefficient *ρ*_*s*_ = 0.79), whereas glutamic acid Cl ranked highest for *N*_max_ (*ρ*_*s*_ = 0.83). lucose, glutamic acid Cl and alanine showed the three strongest positive associations with both parameters, although their order differed (Fig. 2E). Because several compound concentrations covaried, permutation importance was interpreted as a measure of model reliance rather than as the contribution of individual nutrients (Fig. 2F; Methods).

Together, these case studies show how `GUIbiont` supports the progression from inspecting individual growth curves to identifying phenotypic patterns across thousands of gene-level profiles or media. Browser-based fitting and clustering can be reproduced and extended through exported Julia scripts, connecting rapid no-code exploration to customizable analysis in `Kinbiont.j1`. `GUIbiont` is MIT-licensed and available at https://github.com/pinheiroGroup/GUIbiont, with source code, documentation and example datasets. A container image is published to the it ub Container Registry as ghcr.io/pinheirogroup/GUIbiont.

## Methods

### GUIbiont architecture

`GUIbiont` is a single-page web application implemented in Julia [10]. The backend is built with `Oxygen.j1`, a Julia web framework layered over `HTTP.j1`, and exposes a REST API of 24 endpoints for the different analysis routes, including data ingestion, preprocessing, clustering, single and batch fitting, and machine learning analysis. `Oxygen.j1` auto-generates an OpenAPI Swagger specification from the route definitions and typed request schemas, providing interactive documentation at `/docs`. All fitting operations and clustering procedures delegate to `Kinbiont.j1` [7]. Downstream analyses use routines from `DecisionTree.j1` [11] and `t ts se.j1` [12], implemented locally in the GUI application. The frontend is a no-framework single-page application, built with HTML, CSS and JavaScript, that communicates with the backend via JSON. Visualizations are rendered client-side using `P1ot1y.js`; the code export editor uses `CodeMirror`.

The interface provides six analysis tabs: (i) *Clean Data* for preprocessing raw plate-reader files; (ii) *Plot Growth* for interactive inspection of individual and replicate-averaged curves; (iii) *Clustering* for grouping growth trajectories; (iv) *Fit Curve* for single-curve or replicate-level parameter inference; (v) *Batch Fit* for applying single or multiple fitting methods to selected curves; and (vi) *Downstream Analysis* for relating fitted parameters to feature matrices using Spearman rank correlations, random forests and partial-dependence analysis.

Analyses performed in the *Fit Curve, Batch Fit* and *Clustering* tabs can be exported from the browser as executable Julia scripts. Each script contains a placeholder for the local source-data path and records the selected wells or experiments, preprocessing settings and `Kinbiont.j1` function calls required to rerun the workflow outside the browser. For representative single-curve and batch-fitting analyses using data from the two case studies, all numerical outputs compared between the browser and the exported scripts were identical (Supplementary Table S2).

### Data ingestion

The *Clean Data* tab includes parsers for BioTek Synergy 1 CSV exports and Tecan Spark XLSX exports and supports plate-layout annotations for 6-, 12-, 24-, 48- and 96-well plates. Each parser converts the detected measurement channels into standardized columnar CSV files, `data_channe1_N.csv`, containing time in hours followed by one column per well. These files provide a common input format for all subsequent analyses. The plate-reader format can be specified by the user or detected from the input file. Reader-specific parsing is implemented through the `P1ateReaderFormat` interface, allowing support for additional instruments through new format adapters.

### Curve inspection

The *Plot Growth* tab allows users to inspect raw OD curves from selected wells, including wells from multiple experiments. It also reports the saturation OD and the maximum specific growth rate computed by `Kinbiont.j1`, as well as the area under the curve calculated by trapezoidal integration. Wells sharing the same condition, antibiotic and measurement channel can be viewed as a replicate group, with individual curves shown together with their mean.

### Single-curve fitting

The *Fit Curve* tab estimates model parameters for an individual well or a replicate-averaged trajectory using models provided by `Kinbiont.j1` [7]. Parametric models are fitted by nonlinear optimization with a relative-error loss function.

#### Background correction

For individual wells, blank subtraction is optional and uses wells annotated as blanks. When no wells carry a blank annotation, `GUIbiont` proposes candidate blank wells detected from the data, using the default flatness and low-OD criteria described for the clustering pre-screen; wells the user accepts are then used in place of the annotation for both single-curve and batch fitting, and the wells actually used are reported alongside the fit. In point-by-point mode, the mean blank trajectory is subtracted at each measurement time. In the two global modes, a single mean calculated across all blank wells and measurement times is subtracted from the complete trajectory. Following point-by-point subtraction or global subtraction in *Shift minimum* mode, a uniform offset *δ* = max {10^−4^ − min_*t*_ *y*_corr_(*t*), 0} is added when necessary, placing the minimum corrected value at {10^−4^. In *Cli to oor* mode, values below 10^−4^ following global subtraction are instead replaced individually with 10^−4^. This floor ensures positive values for relative-error fitting and is negligible relative to the uncertainty of the OD measurements. When blank subtraction is not applied, parametric fitting instead replaces values below 0.01 with 0.01, because the relative-error loss divides by the predicted values and is therefore sensitive to near-zero observations.

#### Replicate averaging

For replicate-averaged fitting, the global blank mean is calculated separately for each source experiment and subtracted from the corresponding curves when annotated blank wells are available. Values below 0.01 are then set to 0.01. The resulting trajectories are truncated to the length of the shortest selected series and averaged by measurement index using the time vector of the first curve. This procedure therefore assumes that all selected curves were acquired using the same measurement schedule.

#### Model fitting and selection

sers select a single growth model and can either run one optimizer or compare several. Each deterministic optimizer is run once, whereas stochastic optimizers are repeated for a user-defined number of runs. Among successful attempts, `GUIbiont` retains the result with the lowest root-mean-square error against the preprocessed observations and overlays the fitted trajectory on the data. The interface reports the selected model, the optimizer used, the fitted parameters, the AICc and the fitting error.

#### Direct estimation of common growth descriptors

For rapid estimation of commonly used growth descriptors, `GUIbiont` provides a log-linear sliding-window procedure that does not require selecting a parametric growth model. The procedure returns *µ*_max_, from the slope of log OD, lag from the tangent intercept, and *N*_max_ using the standard empirical procedure implemented in `Kinbiont.j1`, as described below in its dedicated section. Users can adjust the smoothing and window settings to reduce sensitivity to measurement noise.

#### Clustering pipeline

As part of this work, we extended the `Kinbiont.j1` framework [7] with methods for clustering complete growth trajectories and made them accessible through the *Clustering* tab in `GUIbiont`. `GUIbiont` assembles the selected curves, handles optional blank correction and presents cluster assignments and diagnostic measures. Per-curve normalization, optional separation of non-growing trajectories and clustering are delegated to `Kinbiont.j1`. The interface supports *k*-means, *k*-medoids, hierarchical clustering and DBSCAN; the analyses reported here used *k*-means.

#### Clustering input

sers can cluster either an uploaded matrix-format CSV or one or more experiments previously imported into `GUIbiont`. In an uploaded CSV, the first data column contains the sampling times and each subsequent column contains one OD trajectory, identified by its column header. For loaded experiments, `GUIbiont` reads the first measurement channel and labels each trajectory by experiment and well. Wells annotated as blanks are excluded from the clustering input and retained separately for optional blank subtraction.

Signal preprocessing. Before clustering, users can optionally apply the blank-correction procedures described for single-curve fitting and smooth the growth trajectories. Blank correction is performed separately within each loaded experiment, so that sample curves are corrected only using blank wells from the same experiment; an uploaded matrix is treated as a single experiment. When annotated blanks are unavailable, users can instead derive a blank signal from candidate non-growing curves identified by the quantile-based pre-screen, the trend-based procedure or both. The selected curves are removed from the clustering input and averaged pointwise to reconstruct the blank trajectory.

Blank correction is followed by optional smoothing of each trajectory. The interface provides LOWESS smoothing [13], selected by default with a bandwidth fraction of 0.05; Gaussian kernel smoothing, with a default bandwidth of twice the median sampling interval; a seven-point rolling average; or no smoothing. Smoothing is applied before the constant-curve pre-screen and z-score normalization.

#### Constant-curve pre-screening

The optional constant-curve pre-screen isolates candidate non-growing trajectories that show little change relative to their baseline before clustering by shape. The criterion is evaluated after any selected blank correction and smoothing. For user-selected quantiles *q*_*ℓ*_ and *q*_*h*_, where *q*_*p*_(*x*_*i*_) denotes the *p*-th empirical quantile of trajectory *i* over time, curve *i* is classified as constant when

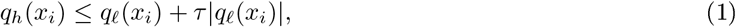

where *τ* is a user-defined tolerance, or when the raw quantile range is negligible,

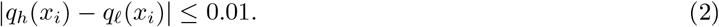

The default settings are *τ* = 0.5, *q*_*ℓ*_ = 0.05 and *q*_*h*_ = 0.95. For clustering methods that use a specified number of groups, all curves meeting the criterion are assigned to a dedicated sentinel cluster with label *k*, and the remaining trajectories are partitioned into *k* − 1 groups. If no curve meets the pre-screen criterion, all *k* groups remain available for clustering.

#### Trend-based separation of flat trajectories

`GUIbiont` can also use a linear trend test to identify trajectories with no detectable temporal trend. A curve is classified as flat when the slope of OD against time is not significantly different from zero in a two-sided *t*-test (default *P* ≥ 0.05). The trend test and constant-curve criterion can be applied independently or together; when both are selected, curves meeting either criterion enter the same non-growing class. For clustering methods with a specified *k*, the remaining trajectories are partitioned into *k* − 1 groups. This option is useful when flatness is better characterized by the absence of an overall temporal trend than by signal change relative to baseline.

After a non-growing group has been identified by the pre-screen or the trend test, users can refine it through the *Nearest curves* view. The interface ranks curves from other clusters by their mean squared pointwise distance from the non-growing centroid, using the selected raw-OD or z-score representation, and displays a user-defined number of candidates. After inspecting a candidate, the user can reassign it to the non-growing class, automatically updating the memberships, centroids and quality metrics.

#### Z-score normalization

Before clustering by trajectory shape, each trajectory is standardized independently across its time points:

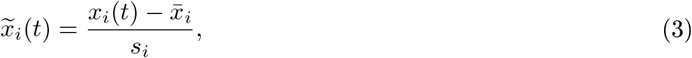

where 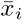 and *s*_*i*_ are the mean and sample standard deviation of curve *i*, respectively. Curves with *s*_*i*_ *<* 10^−12^ are mapped to the zero vector. This transformation removes differences in baseline OD and signal amplitude.Thus, cluster assignments reflect standardized temporal profiles, whereas the original OD-scale can be used to show the average signal magnitude and variation within each resulting group.

#### Clustering algorithms and centroids. GUIbiont

supports *k*-means, *k*-medoids, hierarchical clustering and DBSCAN, all applied to the z-score-normalized trajectories. For *k*-means, `GUIbiont` s default method, three initializations are performed by default using a fixed pseudorandom seed, and the partition with the lowest within-cluster sum of squares (WCSS) is retained. Users can adjust the number of initializations, the maximum number of iterations and the convergence tolerance. *k*-medoids operates on pairwise Euclidean distances between trajectories. Hierarchical clustering uses Ward linkage by default, with average, complete and single linkage also available, and the resulting dendrogram is cut to form the user-requested number of groups *k*. For these three methods, *k* can be chosen using the diagnostics described in the next section. DBSCAN instead does not receive *k* as input and automatically identifies density-connected groups from a user-defined neighborhood radius and minimum neighborhood size; trajectories outside the resulting groups are labeled as noise, while optional non-growing trajectories also sit in their own cluster.

For each cluster *c*, `GUIbiont` computes a new centroid in normalized space,

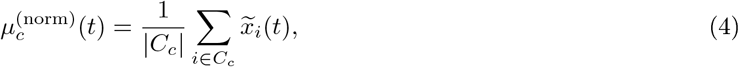

and a corresponding centroid on the pre-normalization OD scale,

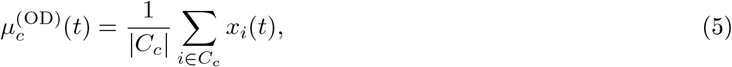

where *C*_*c*_ is the set of trajectories assigned to cluster *c* after cluster assignment. The browser displays the OD-scale centroid and pointwise standard deviation by default, with the normalized representation available as an alternative. This pointwise-mean centroid is computed identically after assignment for all four algorithms and should not be confused with each algorithm s internal representative.

#### Elbow support for choosin

*k*. For clustering methods that require the number of clusters to be chosen in advance, the interface can repeat the analysis over candidate cluster counts *k* ∈ {1, …, *k*_max_}, with *k*_max_ = 10 by default. If the non-growing sentinel is enabled and populated, it is included in the reported number of clusters. Each candidate partition uses the same time-alignment, smoothing and flat-curve settings as the final analysis. For *k* = 1, the non-growing pre-screen is disabled so that WCSS provides the single-cluster baseline. The selected blank correction is also applied consistently during the sweep. The sweep is unavailable for DBSCAN, which does not use *k*.

For each candidate partition, WCSS is calculated as the sum of squared Euclidean distances between each standardized trajectory and the centroid of its assigned cluster. Trajectories assigned to the non-growing class and DBSCAN noise points are excluded from this calculation. When at least three candidate values are evaluated, the interface reports an elbow suggestion, defined as the value *k* that maximizes the second finite difference of the WCSS curve:

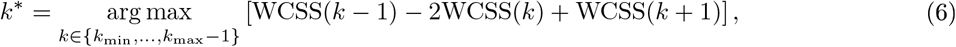

where *k*_min_ = 3 when the non-growing sentinel is populated and *k*_min_ = 2 otherwise. Because the non-growing sentinel cluster only exists once *k* ≥ 2, WCSS(1) is always computed on the full dataset while WCSS(*k* ≥ 2) excludes the sentinel whenever the pre-screen populates it - so the two are not comparable. Thus, *k* = 1 is dropped and the maximization starts from *k* = 3, since the resulting drop from WCSS(1) to WCSS(2) would otherwise reflect the smaller growing population left after removing non-growing trajectories, not a better shape split. With *k*_max_ ≤ 3, this leaves no valid candidate and the search falls back to the original case with *k*_min_ = 2.

The elbow is a diagnostic aid rather than an automatic final decision; the user retains the final choice of *k*. The interface also reports the mean silhouette, Calinski-arabasz, Xie-Beni, Davies-Bouldin and Dunn indices, computed using `C1ustering.j1` [14-19], which characterize candidate partitions and provide alternative suggestions. Quality indices are calculated in normalized trajectory space, with DBSCAN noise points excluded. For datasets containing more than 5,000 trajectories, silhouette values and the Dunn index are skipped, as they require computationally expensive calculations on full pairwise-distance matrices.

### Batch fitting and model selection

`GUIbiont` applies `Kinbiont.j1` [7] in parallel to selected nonblank wells from a previously imported experiment. sers can fit a single model or compare multiple models from the `Kinbiont.j1` model library, which currently contains over 30 parametric and ODE-based growth models.

When multiple candidate models are fitted, `GUIbiont` compares them using the corrected Akaike information criterion (AICc)

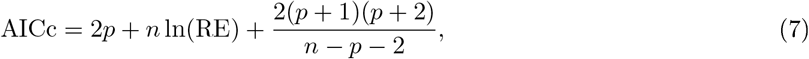

where *p* is the number of fitted model parameters, *n* is the number of observations retained for fitting, and the relative error (RE) is

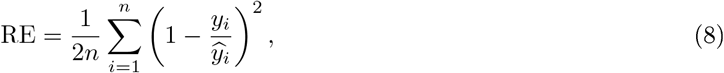

with *y*_*i*_ and 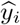 the observed and model-predicted OD values. The model with the smallest AICc value is retained.

Every optimizer receives the same preprocessing options and the same preprocessed curve described for single-curve fitting above. By default, curves with an OD range below 0.02 are excluded from parametric fitting as candidate flat trajectories; this threshold can be adjusted or disabled. Successful fits are returned in a downloadable CSV containing the selected model, its fitted parameters, AICc and fitting-error measures.

### Log-linear parameter estimation

#### Log-linear sliding-window *µ*_max_ estimation

As a model-free alternative to parametric growth models, `GUIbiont` provides a log-linear estimator of the maximum specific growth rate. After any selected blank correction and negative-value handling, the OD values supplied to the logarithmic transformations are constrained to remain positive by the 10^−4^ floor, which is applied whether or not blank subtraction was selected. The log-linear route therefore retains low-OD signal that the 0.01 floor of parametric fitting would remove; the same 10^−4^ floor is used for the log-linear values reported alongside parametric batch fits, so that they match a standalone log-linear analysis of the same curves. By default, each curve is smoothed using a rolling average of width *w*_avg_ (default 7 points). Local specific growth rates are then estimated by ordinary least-squares (OLS) regression,

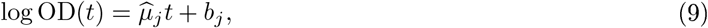

over successive windows of width *w*_deriv_ (default 7 points).

For the default `type_of_win = “maximum”` setting, the routine identifies the largest local slope and searches outward from its position to define a candidate exponential interval. The lower slope threshold is the *α*-quantile of the local-slope distribution, where *α* is specified by `thresho1d_of_exp` (default 0.9).

In each direction, the search stops when two consecutive local slopes below this threshold are encountered. A single local slope below the threshold does not terminate the search and may therefore remain within the selected interval; the search stops only after two consecutive sub-threshold slopes are encountered. If the candidate interval is shorter than a minimum size controlled by *w*_min_ (default 7 points), it is replaced by a window centered on the maximum-slope position. If the centered window extends beyond the smoothed trajectory, its start or end indices are limited to the available observations.

A final OLS regression of log-transformed OD against time is fitted over the selected interval. Its final slope is reported as *µ*_max_, together with its standard error, the interval bounds *t*_start_ and *t*_end_, the largest slope obtained from the preliminary sliding-window regressions, the doubling time, computed as ln(2)*/µ*_max_ and the cofficient of determination *R*^2^, obtained over the final interval. Because the interval is selected from the upper tail of the local-slope distribution around its maximum, the procedure tends to exclude stationary and declining portions of the trajectory without applying a separate stationary-phase truncation before the log-linear fit.

The I fitting routes require at least max {10, *w*_deriv_ + *w*_min_ + 2} valid measurements. This corresponds to 16 measurements under the default settings. In batch analyses, curves whose range max(OD) − min(OD) is smaller than a user-selected threshold are skipped; the default threshold is 0.02. Curves with insufficient data, and curves whose fits raise an exception, are recorded as failed, whereas curves excluded by the flat-signal threshold are recorded as skipped. A candidate fit is accepted only if the estimated growth rate exceeds 10^−6^ h^−1^ and the corresponding *R*^2^ is finite; this excludes growth-rate estimates arising from floating-point noise on flat curves. A fit that completes without returning valid log-linear parameters is retained with `1og1in_converged = fa1se`. These cases do not interrupt the analysis of the remaining curves.

#### Model-free lag and estimated saturation OD

The selected log-linear fit is also used to derive model-free estimates of the lag time and saturation OD:

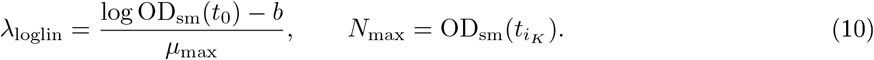

ere, OD_sm_(*t*) denotes the positive smoothed OD trajectory, *b* and *µ*_max_ are the intercept and slope of the final log-linear regression, and *i*_*K*_ is the final stationary-phase cutoff index defined below.

The lag estimate is obtained by extrapolating the fitted exponential trajectory back to the initial smoothed OD [20]. It is defined when both OD_sm_(*t*_0_) *>* 0 and *µ*_max_ *>* 0; otherwise, no finite value is returned.

To estimate *N*_max_, observations satisfying OD_sm_ *>* 0.02 are retained. Local specific growth rates are re-computed on the trajectory using sliding-window OLS regressions with the same width *w*_deriv_ used by the log-linear method. The stationary-phase threshold is defined heuristically as *g*_thr_ = 0.05 × *µ*_max_. Starting from the position of the largest recomputed local slope, the algorithm searches for the first sequence of five consecutive local slopes below *g*_thr_. This position defines a candidate stationary-phase cutoff.

To account for a possible delay between the decline in growth rate and the observed OD plateau, the final cutoff is placed at the largest smoothed OD between the candidate cutoff and the following five observations. Stationary-phase detection is performed only when more than five local slopes exceed *g*_thr_. If this condition is not satisfied, or if no sequence of five consecutive sub-threshold slopes is found, the final observation is used as the cutoff.

For each successful log-linear fit, *λ*_loglin_ and *N*_max_ are returned alongside *µ*_max_ and can be used as targets in subsequent downstream analyses.

### Downstream machine learning

`GUIbiont` relates fitted kinetic parameters to user-supplied experimental features using Spearman rank correlations and random-forest regression implemented with local Julia routines. Users upload a batch-fit results CSV and a feature-matrix CSV. A label column is selected from the batch-fit table, while the first column of the feature matrix contains the corresponding labels. Rows are matched between the two files by label. For each selected fit parameter, `GUIbiont` trains a random-forest regressor containing 100 trees with a maximum depth of 5. At each split, round 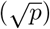 of the *p* predictors are considered. Each tree is fitted to a bootstrap sample containing 70% of the complete observations, drawn with replacement. The minimum leaf and split− sizes are 5 and 2, respectively, and a fixed pseudorandom seed of 42 is used. Predictive performance is evaluated by fivefold cross-validation using the forest settings and summarized by the coefficient of determination *R*^2^. `GUIbiont` clips each fold s *R*^2^ to the interval [− 1, 1] before computing the displayed mean and standard deviation.

#### Spearman rank correlations

For each numeric kinetic parameter and feature, `GUIbiont` calculates the Spearman rank correlation *ρ*_*s*_ across matched rows, excluding missing values, separately for each parameter-feature pair. This measure captures monotonic associations without assuming linearity or normally distributed residuals. For each parameter, the resulting correlations are displayed as a ranked bar chart.

#### Feature importance

Because the target variables are continuous, node impurity is based on squared prediction error. Feature importance is quantified by mean decrease in impurity: the impurity reductions attributed to each feature are normalized within each tree and then averaged across the forest. These rankings summarize how the fitted model uses the available predictors and can capture associations involving nonlinearities and interactions that may not be captured by pairwise correlations.

#### Permutation importance

Impurity-based importance can be difficult to interpret when predictors are correlated. When correlated predictors provide similar reductions in impurity, a tree selects one for the split; the selected predictor receives the associated importance, whereas the other may receive little additional importance. Thus, as a complementary measure, `GUIbiont` calculates permutation importance using `DecisionTree.permutation_importance`. Each predictor is independently shuffled 10 times while the remaining predictors are left unchanged, and the resulting decrease in the fitted forest s training-set *R*^2^ is reported as a mean and standard deviation. This measures the model s reliance on each predictor, but not its unique or causal contribution. Because correlated predictors may compensate for one another after permutation, the resulting importance values are interpreted as descriptive and hypothesis-generating.

#### Partial dependence plots

For the five predictors with the highest impurity-based importance for each kinetic parameter, `GUIbiont` calculates partial-dependence curves over a grid of 30 equally spaced values spanning the predictor s observed range. At each grid value, the selected predictor is assigned that value in every training row, while the remaining predictors retain their observed values; the resulting random-forest predictions are then averaged. These curves summarize the fitted model s marginal response to each predictor. When predictors are correlated, the calculation may evaluate combinations that are uncommon in the data and should therefore be interpreted cautiously.

### Datasets

#### *E. coli* Keio knockout collection

rowth curves were obtained from [8], which assayed 3,909 single-gene deletion strains from the *E. coli* Keio collection, grown in LB and M63 minimal media with three independent biological replicates per strain and medium. The strains represented 3,885 unique gene symbols. Records were grouped by gene symbol and medium and averaged pointwise, yielding 3,885 gene-level trajectories per medium and 7,770 gene-medium trajectories in total. OD measurements were collected at 15-min intervals.

For clustering, a shared set of complete replicate-mean trajectories was prepared for the constant-curve pre-screen and for trajectory clustering because replicate recordings ended at different times. Dataset-specific preparation is described in the Supplementary Material. The constant-curve criterion was evaluated using *q*_*ℓ*_ = 0.05, *q*_*h*_ = 0.95 and *τ* = 0.5. It identified 97 non-growing trajectories in M63 and none in LB. Clustering was performed without blank subtraction or smoothing after applying per-trajectory z-score normalization. WCSS was evaluated for *k* = 1, …, 10, with *k* = 1 providing the single-cluster baseline against which the reduction at larger *k* is read. The automated WCSS elbow diagnostic suggested *k* = 2 for LB and *k* = 3 for M63; these values were used throughout the manuscript. The M63 solution comprised two dynamic clusters and the separate non-growing class.

Enrichment among the 97 M63 non-growing gene-level profiles was evaluated for two KEGG-derived gene sets: amino-acid biosynthesis (`eco01230`) and a broader biosynthesis-related set comprising amino-acid biosynthesis, cofactor biosynthesis, purine metabolism and pyrimidine metabolism (`eco01230, eco01240, eco00230` and `eco00240`) [21]. Both sets were restricted to genes represented in the Keio screen. Enrichment relative to the remaining collection was tested using one-sided Fisher s exact tests. Gene-level pathway memberships and the corresponding contingency-table counts are provided in Supplementary Data S2.

Log-linear batch fitting was run separately for LB and M63 through the `GUIbiont/api/batch-fit-1og1in` endpoint using `b1ank_subtraction = fa1se`, `type_of_smoothing = “ro11ing_avg”`, `pt_avg = pt_smoothing_derivative = pt_min_size_of_win = 5`, `type_of_win = “maximum”`, `thresho1d_of_exp = 0.9` and `skip_f1at_thresho1d = 0`. All other endpoint options were left at their default values. The runtime reported in the main text was measured on a workstation with a six-core Intel Core i7-9850 processor, 48 B RAM, Julia 1.12.6 and Linux 6.8. The analysis scripts and processed results are available at https://github.com/pinheiroGroup/eco1i-knockout-growth-at1as.

#### Chemically defined media screen

Growth curves were obtained from a published screen of wild-type *E. coli* K-12 BW25113 grown in chemically defined media [9]. The dataset comprised seven experimental rounds, 1,029 unique media assembled from 44 compounds and four to twelve replicate cultures for most media, with base-pattern reference media replicated more heavily (up to 588 curves for a single medium). OD_600_ was measured every 30 min for 18 to 48.5 h, yielding 13,608 growth-curve records.

The growth curves were provided in seven Excel files, one for each experimental round, and converted into `GUIbiont` experiment folders using `preprocess.py`. Records containing fewer than ten numeric OD measurements were excluded during conversion. This removed 208 records, including all records for three media, and left 13,400 curves from 1,026 media. The remaining measurements were retained without initial-OD subtraction or missing-value imputation. Log-linear fitting was run separately for each experimental round through the `GUIbiont/api/batch-fit-1og1in` endpoint using `b1ank_subtraction = fa1se`, `type_of_smoothing = “ro11ing_avg”`, `pt_avg = pt_smoothing_derivative = pt_min_size_of_win = 7`, `type_of_win = “maximum”`, `thresho1d_of_exp = 0.9` and `skip_f1at_thresho1d = 0`. All other end-point options were left at their default values. The estimator returned usable fits for 13,349 curves; the remaining 51 did not return a finite positive *µ*_max_.

For downstream analysis, fitted parameters were matched by curve label to the corresponding medium identifier, and replicate estimates from the same medium were averaged. Each of the 1,026 retained media was therefore represented by one parameter row and one matching feature row containing the concentrations of the 44 compounds. Replicate averaging was performed before cross-validation, ensuring that each medium occurred in only one fold. The parameter and feature matrices were submitted to the `GUIbiont/api/m1-downstream` endpoint with *µ*_max_ and *N*_max_ as targets. Tangent-intercept lag was retained in the complete fitting output but was not used as a downstream target. Spearman correlations, random-forest cross-validation and permutation importance were calculated using the procedures described in the *Down-stream machine learning* section. Complete per-curve fitting results are provided in Supplementary Data S3, and the downstream input matrices and outputs are provided in Supplementary Data S4. Analysis scripts are available at https://github.com/pinheiroGroup/chemica1-media-ana1ysis.

## Supporting information

Supplementary Data S2 (KEGG pathway memberships)

Supplementary Data S2 (non-growing enrichment)

Supplementary Data S4 (compound matrix)

Supplementary Data S4 (cross-validation)

Supplementary Data S4 (downstream results)

Supplementary Data S4 (parameter matrix)

## Code availability

`GUIbiont` is released under the MIT license at https://github.com/pinheiroGroup/GUIbiont, with source code, documentation and example datasets. The results reported here were produced with `GUIbiont` v1.1.1 and `Kinbiont.j1` v1.5.1 (DOI: 10.5281/zenodo.21921009), running on Julia 1.12.6. `GUIbiont` commits the exact dependency resolution in `Manifest.tom1`. The versioned container image is built from that manifest without re-resolving dependencies and is published to the it ub Container Registry by the repository s CI workflow as ghcr.io/pinheirogroup/GUIbiont:v1.1.1. Analysis scripts for the two case studies are available at https://github.com/pinheiroGroup/eco1i-knockout-growth-at1as and https://github.com/pinheiroGroup/chemica1-media-ana1ysis. Each repository is tagged at the revision used for this manuscript (`v1.0.8` for the *E. coli* Keio knockout atlas and `v1.0.8` for the chemical-media analysis), and `GUIbiont` v1.1.1 is archived at 10.5281/zenodo.21930894.

The comparison between browser results and exported scripts reported in Supplementary Table S2 is produced by an automated harness distributed with the manuscript sources, which records the repository revisions, Julia version and platform of each run alongside the per-quantity comparison.

## Data availability

The *E. coli* Keio collection growth data analyzed here were obtained from [8] and the chemically defined media screen from [9]. Derived per-curve and per-gene results, the matrices submitted to the downstream analysis and the enrichment tables are provided as Supplementary Data S1-S4.

## Acknowledgements

We thank Fabrizio Angaroni for contributions during the initial development of `GUIbiont`, and Claudio Del Fatti for extensive user testing and feedback throughout the development of the software. This work was funded by uman Technopole.

## Author contributions statement

Conceptualization: F.P., E.Z.A. and E.O. Methodology: E.Z.A., E.O. and F.P. Software: E.Z.A. and E.O. Validation: E.Z.A. and E.O. Formal analysis: E.Z.A., E.O. and F.P. Visualization: E.Z.A., E.O. and F.P. Writing - original draft: E.Z.A., E.O. and F.P. Writing - review & editing: E.Z.A., E.O. and F.P. Supervision: F.P. Funding acquisition: F.P.

## References

1. Kahm, M., Hasenbrink, H., Lichtenberg-Fraté, H., Ludwig, J. & Kschischo, M. grofit: fitting biological growth curves with R. Journal of statistical software 33, 1–21 (2010).

2. Sprouffske, K. & Wagner, A. Growthcurver: an R package for obtaining interpretable metrics from microbial growth curves. BMC bioinformatics 17, 172 (2016).

3. Midani, F. S., Collins, J. & Britton, R. A. AMi A: software for automated analysis of microbial growth assays. Msystems 6, e00508–21 (2021).

4. Wirth, N. T., Funk, J., Donati, S. & Nikel, P. I. QurvE: user-friendly software for the analysis of biological growth and fluorescence data. Nature Protocols 18, 2401–2403 (2023).

5. Blazanin, M. gcplyr: an R package for microbial growth curve data analysis. BMC bioinformatics 25, 232 (2024).

6. Bradley, S. A., Webel, H., Donati, S. & Acevedo-Rocha, C. G. growthcurves: A platform for human-in-the-loop analysis of biological growth curves. bioRxiv) 2026–05 (2026).

7. Angaroni, F., Peruzzi, A., Alvarenga, E. Z. & Pinheiro, F. Translating microbial kinetics into quantitative responses and testable hypotheses using Kinbiont. Nature Communications 16, 6440 (2025).

8. Lao, Z. & Ying, B.-W. Growth dynamics of 3,909 Escherichia coli single-gene knockouts in rich and minimal media. Scientific Data 13, 717 (2026).

9. Aida, H. & Ying, B.-W. Population dynamics of Escherichia coli growing under chemically defined media. Scientific Data 12, 984 (2025).

10. Bezanson, J., Edelman, A., Karpinski, S. & Shah, V. B. Julia: A fresh approach to numerical computing. SIAM Review 59, 65–98 (2017).

11. Sadeghi, B. et al. DecisionTree.jl: A Julia im lementation of the CART decision-tree and random-forest algorithms version 0.12.4. Oct. 2023. https://github.com/JuliaAI/DecisionTree.jl/releases/ta438g/v0.12.4

12. JuliaStats contributors. StatsBase.jl: Basic statistics for Julia version 0.34.12. 2026. https://github.com/JuliaStats/StatsBase.jl/releases/tag/v0.34.12.

13. Cleveland, W. S. & Grosse, E. Computational methods for local regression. Statistics and Computing 1, 47–62 (1991).

14. JuliaStats contributors. Clustering.jl: Methods for data clustering and evaluation of clustering quality version 0.15.8. 2025. https://github.com/JuliaStats/Clustering.jl/releases/tag/v0.15.8.

15. Rousseeuw, P. J. Silhouettes: A graphical aid to the interpretation and validation of cluster analysis. Journal of Com utational and A lied Mathematics 20, 53–65 (1987).

16. Calinski, T. & Harabasz, J. A dendrite method for cluster analysis. Communications in Statistics 3, 1–27 (1974).

17. Xie, X. & Beni, G. A validity measure for fuzzy clustering. IEEE Transactions on Pattern Analysis and Machine Intelligence 13, 841–847 (1991).

18. Davies, D. L. & Bouldin, D. W. A cluster separation measure. IEEE Transactions on Pattern Analysis and Machine Intelligence PAMI-1, 224–227 (1979).

19. Dunn, J. C. Well-separated clusters and optimal fuzzy partitions. Journal of Cybernetics 4, 95–104 (1974).

20. Zwietering, M. H., Jongenburger, I., Rombouts, F. M. & Van t Riet, K. Modeling of the bacterial growth curve. A lied and environmental microbiology 56, 1875–1881 (1990).

21. Kanehisa, M., Furumichi, M., Sato, Y., Matsuura, Y. & Ishiguro-Watanabe, M. KEGG: biological systems database as a model of the real world. Nucleic acids research 53, D672–D677 (2025).

