## Supplementary Material for "No-code microbial growth phenotyping with GUIbiont"

### Contents

1 **Comparison with existing tools** 2

2 **Reproducibility: GUI vs. exported Julia script** 2

3 **Extended results: *E. coli* Keio knockout collection** 3

4 **Extended results: chemically defined media screen** 4

5 **Supplementary Data** 5

[†]These authors contributed equally to this work

### 1 Comparison with existing tools

Table S1 compares `GUIbiont` with a selected set of established microbial growth-curve analysis tools. The comparison focuses on interfaces and analytical workflows rather than modeling breadth: AMiGA provides Gaussian-process inference, `Kinbiont.jl` includes symbolic regression, and QurvE supports dose-response analysis. Built on `Kinbiont.jl`, `GUIbiont` combines a local browser interface for no-code analysis with phenotypic clustering, metadata-linked analysis, a REST API and executable Julia-script export. `Kinbiont.jl` was expanded in parallel during the development of `GUIbiont`, including the clustering and non-growing pre-screen introduced in this work. Accessing these capabilities directly through `Kinbiont.jl` requires Julia programming, whereas `GUIbiont` exposes them through browser-based workflows.

| Feature | GUIbiont | Kinbiont.jl | grofit | Growth-curver | AMiGA | QurvE | gcplyr |
|---|---|---|---|---|---|---|---|
| **A. Tool identity** | | | | | | | |
| **Language / runtime** | Julia + JS | Julia | R | R | Python | R | R |
| **Interface** | Browser GUI + REST API | Julia REPL/script | R script | R script | CLI | R script + Shiny | R script |
| **Latest release** | v1.1.1 (2026-08) | v1.5.1 (2026-08) | v1.1.1-1 (2014) archived 2018 | v0.3.1 (2020-10) | v3.0.4 (2025-03) | v1.1.2 (2025-09) | v1.12.0 (2025-07) |
| **License** | MIT | MIT | GPL-2 (legacy) | GPL $\geq 2$ | GPL-3.0 | GPL $\geq 3$ | MIT |
| **B. Interface and reproducibility** | | | | | | | |
| **No-code, no install required** | ✓ | ✗ | ✗ | ✗ | ✗ | ❍ | ✗ |
| **Script export from GUI (reproducibility bridge)** | ✓ | – | – | – | – | ✗ | – |
| **REST / HTTP API** | ✓ | ✗ | ✗ | ✗ | ✗ | ✗ | ✗ |
| **C. Curve fitting and model selection** | | | | | | | |
| **Parametric models (logistic, Gompertz, Baranyi, Richards, aHPM)** | ✓ | ✓ | ❍ | ❍ | ✗ | ❍ | ✗ |
| **Non-parametric / model-free fit (log-linear, spline, GP)** | ✓ | ✓ | ❍ | ✗ | ✓ | ❍ | ✓ |
| **Automatic model selection (AIC / AICc / BIC)** | ✓ | ✓ | ✓ | – | – | ✓ | – |
| **Batch fitting (whole-plate / multi-plate)** | ✓ | ✓ | ✓ | ✓ | ✓ | ✓ | ✓ |
| **D. Phenotypic analysis and downstream ML** | | | | | | | |
| **Growth-curve clustering and non-growing pre-screen** | ✓ | ✓ | ✗ | ✗ | ✗ | ✗ | ✗ |
| **Metadata-linked downstream analysis** | ✓ | ✓ | ✗ | ✗ | ❍ | ❍ | ❍ |
| **E. Data ingestion** | | | | | | | |
| **Vendor plate-reader format parsers** | ✓ | ❍ | ❍ | ❍ | ✓ | ✓ | ✓ |
| **Plate-layout (96/48/6-well) annotation parser** | ✓ | ❍ | ✗ | ✗ | ✓ | ✓ | ✓ |

Table S1: **Comparison of microbial growth-curve analysis tools.** ✓ full support; ❍ partial or limited support; ✗ absent; – not applicable. Marks refer to functionality provided directly by each tool. The `GUIbiont` column is highlighted in pale blue. The compared tools are `Kinbiont.jl` [1], grofit [2], Growthcurver [3], AMiGA [4], QurvE [5] and gcplyr [6]. Package versions and features were checked in August 2026.

### 2 Reproducibility: GUI vs. exported Julia script

To verify that analyses exported from `GUIbiont` can be reproduced outside the browser, an automated harness, using release `v1.0.8` of the Keio repository and `v1.0.8` of the chemical-media repository, ran four analyses through the browser interface, recorded the values the interface displayed, downloaded the generated `guibiont_analysis.jl` script through the export dialog and executed it in a separate Julia process. Every interaction used the same code path as a manual session. The downloaded script was preserved unchanged; a verification copy supplied the local data path and recorded its outputs for automated comparison. The single-curve case used the adjusted heterogeneous population model (aHPM), a four-parameter ODE model in which dormant cells transition to an actively growing population [1]; although aHPM was not used in the case studies, its multi-parameter ODE formulation provides a more demanding test of the exported workflow. The four cases were: (i) a single-curve aHPM fit with its log-linear companion, on the Keio *aaeA* gene-level profile in M63; (ii) log-linear batch fitting of one chemical-media round (2,628 curves), using the settings of the chemical-media case study; (iii) parametric batch fitting with the log-linear companion on ten Keio LB gene-level profiles, which exercises the companion columns written to the batch results file; and (iv) clustering of 44 sample curves from a 48-well plate into three groups.

Fitted parameters, error measures and log-linear descriptors were compared per curve rather than only through summary statistics; counts, model names, optimizer choices and clustering diagnostics were also checked. This yielded 26,434 quantities in total. Floating-point values were compared with an absolute tolerance of $10^{-6}$; counts, model names and optimizer choices were required to match exactly. Cluster assignments were compared up to a permutation of cluster labels. Of the 26,434 quantities, 26,197 were bitwise identical, three clustering quality indices agreed to within $2.8 \times 10^{-14}$, and 234 log-linear descriptors were undefined in both the browser and the script, all belonging to the 26 chemical-media curves in the batch round that did not meet the log-linear acceptance threshold. Representative values are listed in Table S2.

| analysis | quantity | GUI value | exported script | absolute difference |
|---|---|---|---|---|
| (i) Keio M63 single-curve aHPM fit on *aaeA* | $\mu$ (gr) | 3.3397733490657084 | 3.3397733490657084 | 0 |
| | exit-lag rate $\alpha$ | 0.001330822690236778 | 0.001330822690236778 | 0 |
| | $K$ ($N_{\max}$) | 1.0526131723073133 | 1.0526131723073133 | 0 |
| | shape $s$ | 0.1059885818132303 | 0.1059885818132303 | 0 |
| | AICc | −438.38333962525115 | −438.38333962525115 | 0 |
| | RMSE | 0.007357501262028934 | 0.007357501262028934 | 0 |
| | log-linear $\mu_{\max}$ | 0.6295254930981222 | 0.6295254930981222 | 0 |
| | log-linear $\lambda$ (h) | 2.553177806751187 | 2.553177806751187 | 0 |
| (ii) Chemical-media `batch-fit-loglin` on bw25113_round01 | converged count | 2,602 | 2,602 | 0 |
| | skipped count | 0 | 0 | 0 |
| | per-curve quantities compared | 26,280 | 26,280 | 0 |
| (iii) Keio LB parametric batch + log-linear companion | curves fitted | 10 | 10 | 0 |
| | per-curve quantities compared | 120 | 120 | 0 |
| (iv) Clustering $k = 3$ | profiles clustered | 44 | 44 | 0 |
| | cluster sizes | 20 / 15 / 9 | 20 / 15 / 9 | 0 |
| | WCSS | 92.59306583569057 | 92.59306583569057 | 0 |
| | mean silhouette | 0.4922662275132982 | 0.49226622751329807 | $1.1\times10^{-16}$ |
| | profiles in a different cluster | 0 | 0 | 0 |

Table S2: **Reproducibility of exported Julia scripts.** Representative numerical outputs obtained through the browser interface and by running the corresponding exported `guibiont_analysis.jl` script. Values are reported at full returned precision. Across all four cases, 26,434 quantities were compared and the largest absolute difference was $2.8 \times 10^{-14}$, in a clustering quality index whose value depends on floating-point summation order. The complete per-quantity comparison, together with the commit identifiers, Julia version and platform of the run that produced it, is provided with the analysis scripts.

### 3 Extended results: *E. coli* Keio knockout collection

#### Trajectory preparation for clustering

The Keio growth-curve dataset analyzed here was obtained from Ref. [7]. Each replicate was extended from its final measurement to 50 h by carrying its final OD forward, and replicates were then averaged pointwise for each gene and medium. The resulting complete 0.25 to 50 h trajectories were used for both the constant-curve pre-screen and shape clustering.

##### Additional enrichment statistics

The amino-acid biosynthesis set contained 103 genes represented in the Keio screen. Among the 97 non-growing gene-level profiles, 49 deletions belonged to this set, compared with 2.57 expected based on its frequency in the complete collection, giving an odds ratio of 70.6. The broader biosynthesis-related set contained 278 genes, with 82 observed among the non-growing profiles compared with 6.94 expected, giving an odds ratio of 100.2. The gene-level pathway memberships are provided in Supplementary Data S2.

##### Additional log-linear results

Estimates of $\mu_{\max}$ were available for 7,767 of the 7,770 gene–medium trajectories. The three unsuccessful M63 trajectories (*leuA*, *lysA* and *nuoB*) returned no finite positive $\mu_{\max}$ and therefore also have undefined tangent-intercept lag $\lambda$, $R^2$ and $N_{\max}$; all missing values are retained in Supplementary Data S1.

#### 4 Extended results: chemically defined media screen

##### Fit completeness across experimental rounds

The chemically defined media growth-curve dataset analyzed here was obtained from Ref. [8]. Quality-control exclusions and usable log-linear fits for each experimental round are summarized in Table S3. The 208 records excluded during data conversion occurred in rounds 4 and 5. Log-linear fitting returned usable fits for all retained records in rounds 6 and 7 and for 13,349 of the 13,400 retained records overall. Records without a usable fit are retained in the complete output together with their fit status.

| experimental round | original records | after quality control | usable fits |
|---|---|---|---|
| Round 1 | 2,628 | 2,628 | 2,602 |
| Round 2 | 2,640 | 2,640 | 2,635 |
| Round 3 | 960 | 960 | 951 |
| Round 4 | 2,640 | 2,574 | 2,567 |
| Round 5 | 2,640 | 2,498 | 2,494 |
| Round 6 | 1,320 | 1,320 | 1,320 |
| Round 7 | 780 | 780 | 780 |
| Total | 13,608 | 13,400 | 13,349 |

Table S3: **Quality control and log-linear fitting across the seven experimental rounds.** Records retained after quality control contained at least ten numeric OD measurements. Usable fits returned a finite positive $\mu_{\max}$.

Supplementary Data S3 contains the complete log-linear output and fit status for all 13,608 original records, including the curve label, medium identifier and fitted parameters. Of these, 13,400 were retained after quality control and 13,349 yielded usable fits. Supplementary Data S4 contains the parameter and compound-concentration matrices for the 1,026 media used in downstream analysis. For each of the 44 compounds, it reports the Spearman correlation with $\mu_{\max}$ and $N_{\max}$, impurity-based random-forest importance and the mean and standard deviation of the permutation importance. Fold-specific and summary cross-validation $R^2$ values are also provided.

### 5 Supplementary Data

The following CSV files accompany this manuscript as Supplementary Data.

| ID | File | Contents |
|---|---|---|
| S1 | `keio_loglin_results.csv` | Per-gene log-linear results for the Keio collection (3,885 genes × 2 media = 7,770 rows). Columns: gene name, Keio JW identifier (`jw_id`), medium (LB / M63), number of replicate curves contributing to the gene-level mean (`n_replicates`; three for most rows and six or nine when multiple source entries mapped to the same gene symbol), $\mu_{\max}$ ($h^{-1}$) with its $1\sigma$ standard error, the largest preliminary sliding-window slope, the exponential-window bounds $t_{\text{start}}$ and $t_{\text{end}}$, doubling time, $R^2$ of the fit on log OD, tangent-intercept lag (h; can be slightly negative when the curve enters exponential phase before the first measurement), $N_{\max}$, and a convergence flag. Lag, $R^2$ and $N_{\max}$ are empty for three trajectories. The log-linear columns were produced by `GUIbiont`'s `/api/batch-fit-loglin` endpoint; `jw_id` and `n_replicates` were joined from metadata generated during gene-level aggregation. |
| S2 | `keio_kegg_pathway_memberships.csv`, `keio_nongrowing_enrichment.csv` | Gene-level KEGG memberships for all 3,885 genes in the screen, flagging the 97 assigned to the non-growing class in M63, membership of the amino-acid biosynthesis set (`eco01230`) and of the wider biosynthesis-related set (`eco01230`, `eco01240`, `eco00230`, `eco00240`), together with the per-pathway flags. The enrichment file gives the complete 2-by-2 contingency table for each set, the observed and expected counts, fold enrichment, odds ratio and the one-sided Fisher exact $P$ value. |
| S3 | `chemical_media_loglin_results.csv` | Log-linear results for every one of the 13,608 chemical-media records, one row each. Columns: experimental round, curve label, medium identifier, fit status, and the log-linear descriptors. The status distinguishes the 208 records excluded during conversion for having fewer than ten numeric OD measurements, the 51 that were fitted but returned no finite positive $\mu_{\max}$, and the 13,349 usable fits; unsuccessful records are retained as rows with empty descriptors. |

| ID | File | Contents |
| --- | --- | --- |
| S4 | chemical_media_parameter_matrix.csv, chemical_media_compound_matrix.csv, chemical_media_downstream_results.csv, chemical_media_cross_validation.csv | Inputs and outputs of the downstream analysis. The two matrices hold the replicate-averaged parameters and the 44 compound concentrations for the 1,026 media, as submitted to /api/ml-downstream. The results file gives, per compound, the Spearman correlation with $\mu_{\max}$ and $N_{\max}$, impurity-based random-forest importance, and the mean and standard deviation of the permutation importance. The cross-validation file reports the fold-level and summary $R^2$ for each target. |

### References


1. Angaroni, F., Peruzzi, A., Alvarenga, E. Z. & Pinheiro, F. Translating microbial kinetics into quantitative responses and testable hypotheses using Kinbiont. *Nature Communications* **16,** 6440 (2025).
2. Kahm, M., Hasenbrink, G., Lichtenberg-Fraté, H., Ludwig, J. & Kschischo, M. grofit: fitting biological growth curves with R. *Journal of statistical software* **33,** 1–21 (2010).
3. Sprouffske, K. & Wagner, A. Growthcurver: an R package for obtaining interpretable metrics from microbial growth curves. *BMC bioinformatics* **17,** 172 (2016).
4. Midani, F. S., Collins, J. & Britton, R. A. AMiGA: software for automated analysis of microbial growth assays. *Msystems* **6,** e00508–21 (2021).
5. Wirth, N. T., Funk, J., Donati, S. & Nikel, P. I. QurvE: user-friendly software for the analysis of biological growth and fluorescence data. *Nature Protocols* **18,** 2401–2403 (2023).
6. Blazanin, M. gcplyr: an R package for microbial growth curve data analysis. *BMC bioinformatics* **25,** 232 (2024).
7. Lao, Z. & Ying, B.-W. Growth dynamics of 3,909 Escherichia coli single-gene knockouts in rich and minimal media. *Scientific Data* **13,** 717 (2026).
8. Aida, H. & Ying, B.-W. Population dynamics of Escherichia coli growing under chemically defined media. *Scientific Data* **12,** 984 (2025).
